# Radiation exposure of the base of the heart accelerates coronary atherosclerosis

**DOI:** 10.1101/2021.04.08.438992

**Authors:** Prerna R. Nepali, Mickael Mathieu, Sarah Kitz, Chiharuko Nakauchi, Karen Gabriels, James Russell, Sebastien Monette, Johannes A. te Poele, Sylvia Heeneman, Andreas Rimner, Yunping Qiu, Irwin J. Kurland, Fiona A. Stewart, Edgar A. Jaimes, Adriana Haimovitz-Friedman

## Abstract

Clinical studies have identified cardiac exposure as an independent predictor for cardiovascular mortality in patients treated with radiation therapy (RT) for thoracic malignancies. Although the mechanisms are not completely understood, the available evidence indicate that direct injury to the coronary arteries endothelium is implicated. In these studies we tested the hypothesis that different areas of the heart are more sensitive to the effects of RT on the formation of atherosclerotic plaque in apolipoprotein E deficient (ApoE^-/-^) mice, a well validated model of atherosclerosis.

**Methods:** ApoE^-/-^ mice on a high fat diet (HFD) received 16Gy cardiac irradiation targeted to the whole or partial (apical or basal) region of the heart at 9 weeks or 16 weeks of age. Atherosclerotic lesions and inflammatory changes in the hearts as compared to control unirradiated mice were assessed eight weeks following radiation.

**Results:** After either basal or whole heart RT at 9 weeks of age the number of subendocardial atherosclerotic lesions at the heart base was higher as compared to unirradiated mice. Irradiation of the apex did not increase the number of subendocardial atherosclerotic lesions in any region. After basal RT at 16 weeks of age the number of coronary and subendocardial atherosclerotic lesions was higher as compared to controls. Neither apical or whole heart RT had an impact on the development or acceleration of lesions in the basal region of the hearts of 16 week old mice, thus demonstrating the adverse impact of basal irradiation. Infiltration of inflammatory cells (CD45^+^ and CD3^+^) and enhanced expression of endothelial adhesion molecules (CD31), were differentially and locally regulated based upon the site of irradiation. In support of a role of eicosanoid mediators for base or whole heart atherogenic irradiation effects, apex irradiation eicosanoid mediators are not clearly atherogenic, in contrast to eicosanoid mediators detected in serum after base heart irradiation. These results indicate that the base of the heart is significantly more prone to the development of atherosclerotic lesions in the coronary arteries post-RT.

**Conclusion:** Our results indicate that the base of the heart is more susuceptible to development of RT-induced atherosclerotic lesions and therefore avoidance from RT direct exposure to this area may reduce the risk for atherosclerotic disease in patients undergoing RT.

## Introduction

Thoracic RT is commonly used in patients with breast cancer [1], Hodgkin Lymphoma (HL) [2], and lung cancer [3]. The benefits of RT are often out-weighed by associated morbidities including increased incidence of CVD as a result of direct irradiation to the heart during RT [4]. CVD is recognized as the leading cause of non-cancer related mortality in survivors of HL [4, 5] and in left-sided breast cancer patients receiving thoracic RT [4]. RT-induced CVD complications include pericarditis, myocardial fibrosis, coronary artery disease, valvular disease, arrhythmias, autonomic dysfunction, vascular calcification and microvasculature injury [4].

Randomized clinical studies have demonstrated reduced rates of recurrence and improved survival in patients that receive RT for early-stage breast cancer. However, long-term follow-up in these studies have shown increased rates of CVD due to incidental exposure of the heart to RT [6]. In a population-based case-control study of major coronary events including myocardial infarction, coronary revascularization or death from ischemic heart disease, it was observed that these events occurred less than 10 years after diagnosis in 44% patients, 10-19 years post diagnosis in 33% patients and 20 or more years later in 23% patients [6]. In addition, reductions in cardiac function have been observed approximately 15 years after RT in HL survivors who received mediastinal RT at a median age of 16.5 years [7]. For 5-year survivors of adolescent or adult HL who received mediastinal RT with a dose equal to or greater than 25Gy, the incidence of heart failure has been shown to markedly increase in a cardiac radiation dose dependent manner [8]. Moreover, patients receiving RT for Stage III non-small-cell lung cancer treated on dose-escalation trials have an association between clinically significant cardiac events and RT doses to the heart [9].

Modern RT regimens with more focused radiation beams, allow tumors to be targeted more precisely and shield the heart and other healthy tissues to minimize the RT-induced injury to normal tissues [10]. Techniques utilized to reduce cardiac radiation exposure include the use of a breast board for patients with breast cancer [11, 12]. Prone positioning is also associated with a reduction in the amount of irradiated lung as well as irradiated cardiac volume in 85% of patients with left breast cancer [13]. However, it remains to be determined whether reductions in radiation exposure to the heart by reducing the RT fields or dose lead to lower incidence of CVD in these patients. In addition, no threshold radiation dose has been reported that can lead to reduced cardiotoxicity [10]. The mechanisms associated with the development of RT-associated CVD are not completely understood. Experimental studies using low doses of total body irradiation in ApoE^-/-^ mice have demonstrated changes in inflammatory markers including CD31, E-selectin, thrombomodulin and vascular cell adhesion molecule-1 (VCAM-1) in the heart at 3 and 6 months post-RT [14]. Injury to the vascular endothelium associated with chronic inflammation resulting in accelerated atherosclerosis are suspected to play a key role in inducing RT-related CVD [15–17].

Eicosanoids are a group of lipid mediators originating from Arachidonic Acid (AA) consisting of prostaglandins (PGs), thromboxanes (TXs), leukotrienes, and lipoxins. The oxidation of AA *via* cyclooxygenase (COX) and lipoxygenase (LOX) activity to produce eicosanoids during inflammation is a key biosynthetic pathway present in all cells. Eicosanoids are involved in blood pressure regulation, renal function, reproduction and host defense against microorganisms and these molecules when generated in excess can induce pain, fever, and diverse inflammatory reaction [18]. COX-1 induced increase in the production of prostacyclin and TX generation is implied in the development of atherosclerosis [19]. COX-1 deletion in ApoE^-/-^ mice and COX-1, COX-2 inhibition in low-density lipoprotein receptor knockout (LDLr^-/-^) mice has been shown to significantly attenuate atherosclerosis lesion development [19, 20].

Epoxyeicosatrienoic acids (EET) and hydroxyeicosatetraenoic acids (HETE) are key metabolites of AA in the cytochrome P450 pathway that are produced by the vascular endothelium [21–23]. EETs have been implied in the regulation of coronary vasomotor tone and myocardial perfusion, inhibition of vascular smooth muscle migration, decreasing inflammation, inhibition of platelet aggregation and decrease adhesion molecule expression, therefore, representing an endogenous protective mechanism against atherosclerosis [22, 23]. 15-LOX enzymatic activity has been shown to have a role in the initiation and development of atherosclerosis [24] (see Overview Fig. 5).

**Figure 1.**
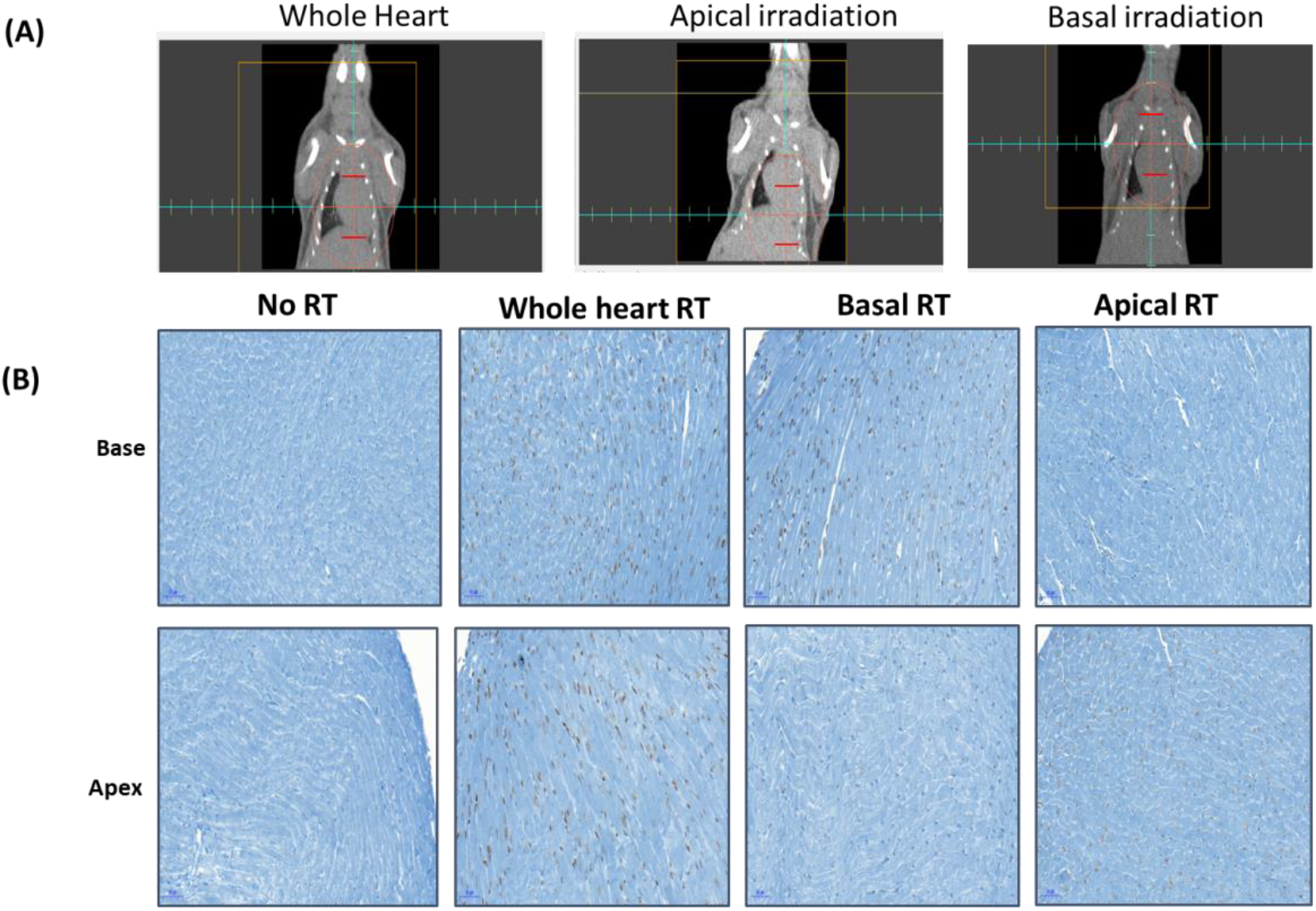
**(A) Irradiation field localization.** Targeted irradiation to specific locations of the heat **(B) Irradiation field localization.** (**i**) γ-H2AX staining after No RT (**ii**) whole heart 16Gy (**iii**) Basal 16Gy (**iv**) Apical 16Gy irradiation

**Figure 2.**
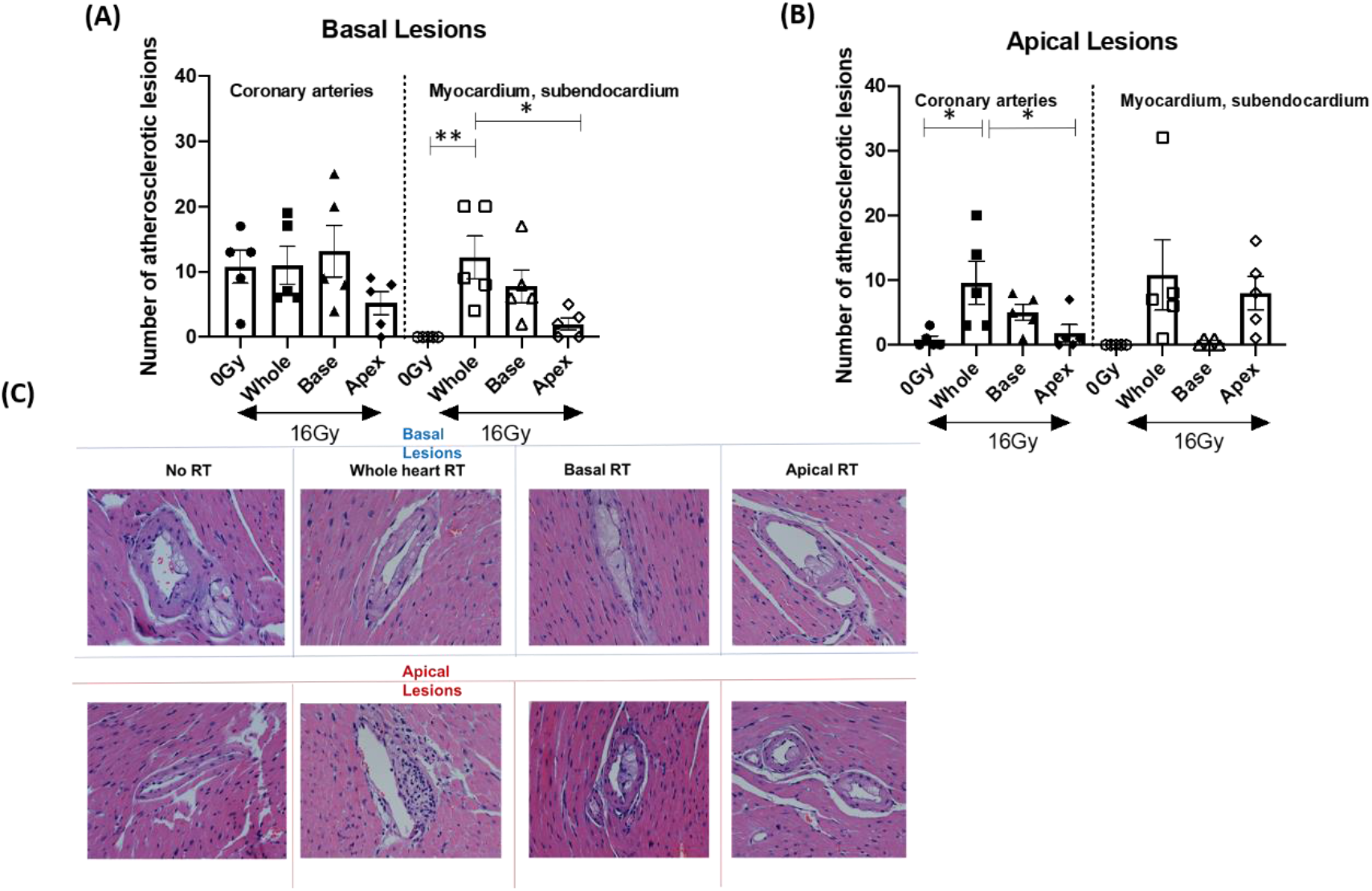
Average number of atherosclerotic lesions 8 weeks post cardiac irradiation of ApoE ^-/-^ mice. **(A)** Basal lesions observed in the coronary arteries or myocardium/subendocardium of mice irradiated with 16 Gy on the whole heart, base or apex, or 0 Gy and **(B)** Apical lesions observed in the coronary arteries or myocardium/subendocardium of mice irradiated with 16 Gy on the whole heart, base or apex, or 0 Gy and. Bars represent mean + SEM. * p<0.05, **p<0.01. n=5 per group, each dot represents number of lesions from the heart of one mouse. **(C)** Representative H&E images of atherosclerotic lesions formed post differential cardiac irradiation.

**Figure 3.**
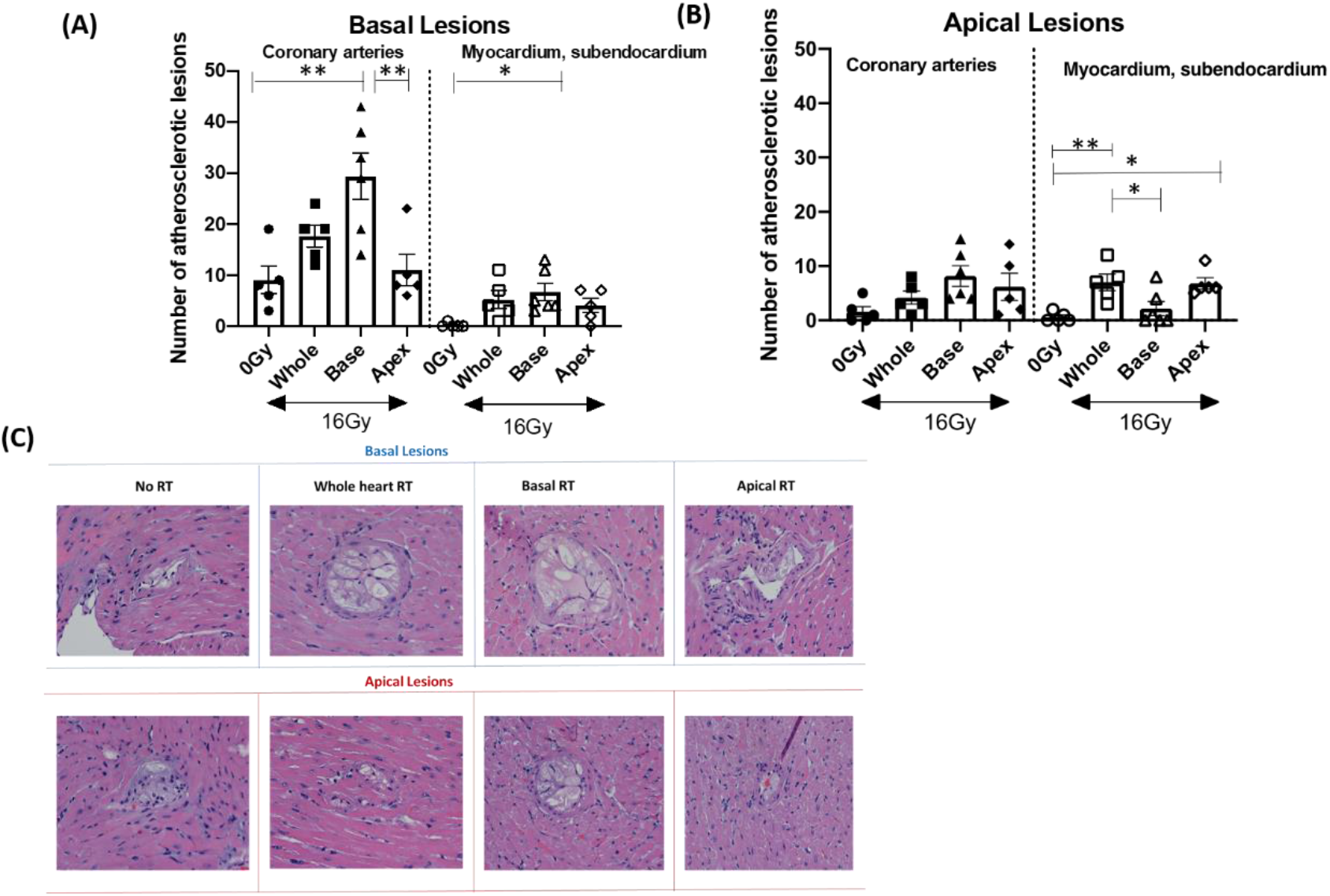
Average number of atherosclerotic lesions 8 weeks post cardiac irradiation of 16 weeks old ApoE ^-/-^ mice. **(A)** Basal lesions observed in the coronary arteries or myocardium/subendocardium of mice irradiated with 16 Gy on the whole heart, base or apex, or 0 Gy and **(B)** Apical lesions observed in the coronary arteries or myocardium/subendocardium of mice irradiated with 16 Gy on the whole heart, base or apex, or 0 Gy and. Bars represent mean ± SEM. * p<0.05, **p<0.01. (C) Representative H&E images of atherosclerotic lesions formed post differential cardiac irradiation.

**Figure 4.**
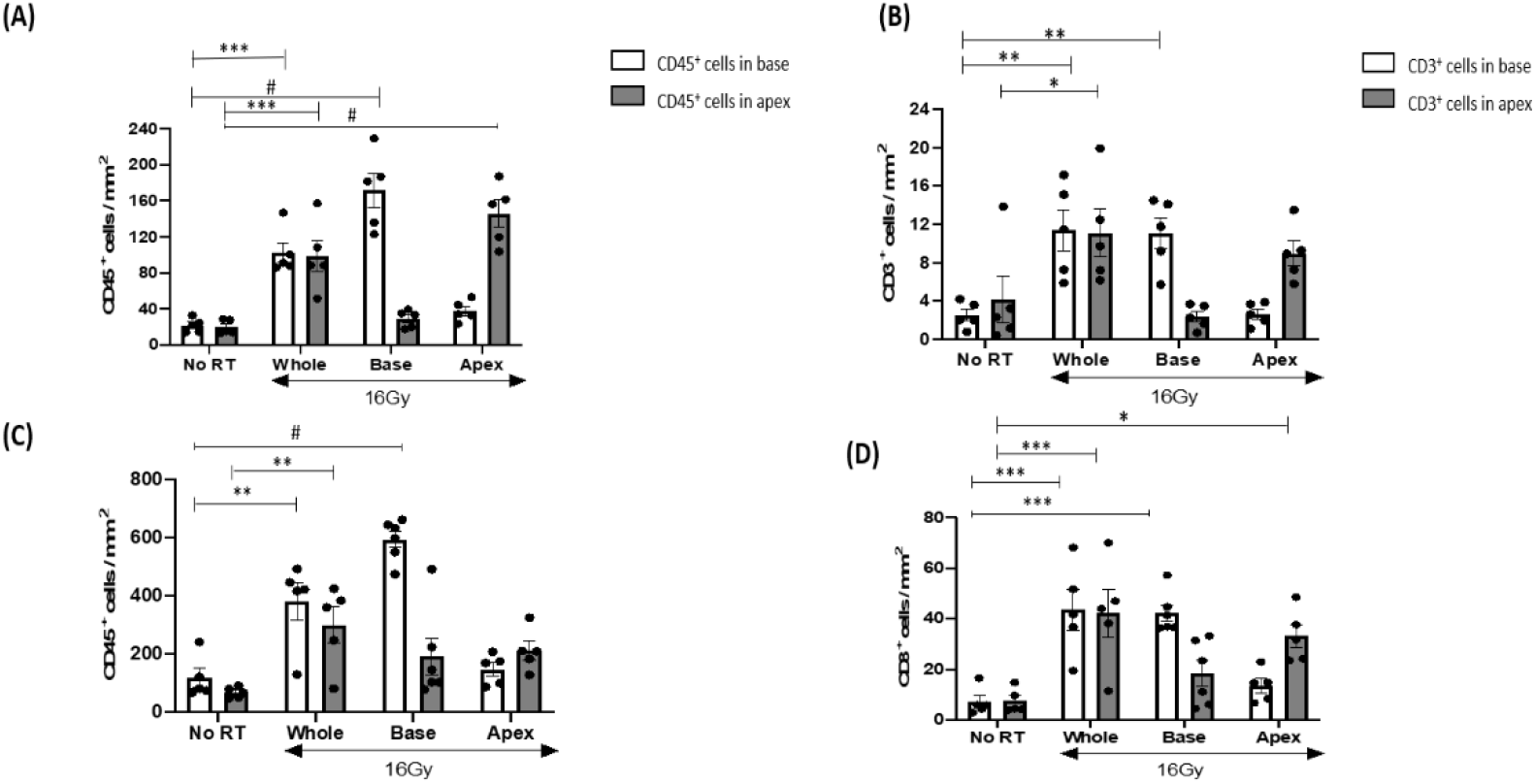
Inflammatory changes observed post differential cardiac irradiation of ApoE^-/-^ mice. CD45^+^cells observed in the base and apex of (A) 9 weeks old and (C) 16 weeks old mice at 8 weeks after whole heart, basal, apical or no irradiation. CD3^+^ cells infiltration in the base and apex of (B) 9 weeks and (D) 16 weeks old mice at 8 weeks after whole heart, basal, apical or no irradiation. Bars represent mean + SEM. * p<0.05, **p<0.01, ***p<0.001, #p<0.0001. n=5 per group, each dot represents number of CD45^+^ or CD3^+^ from the heart of one mouse.

**Figure 5.**
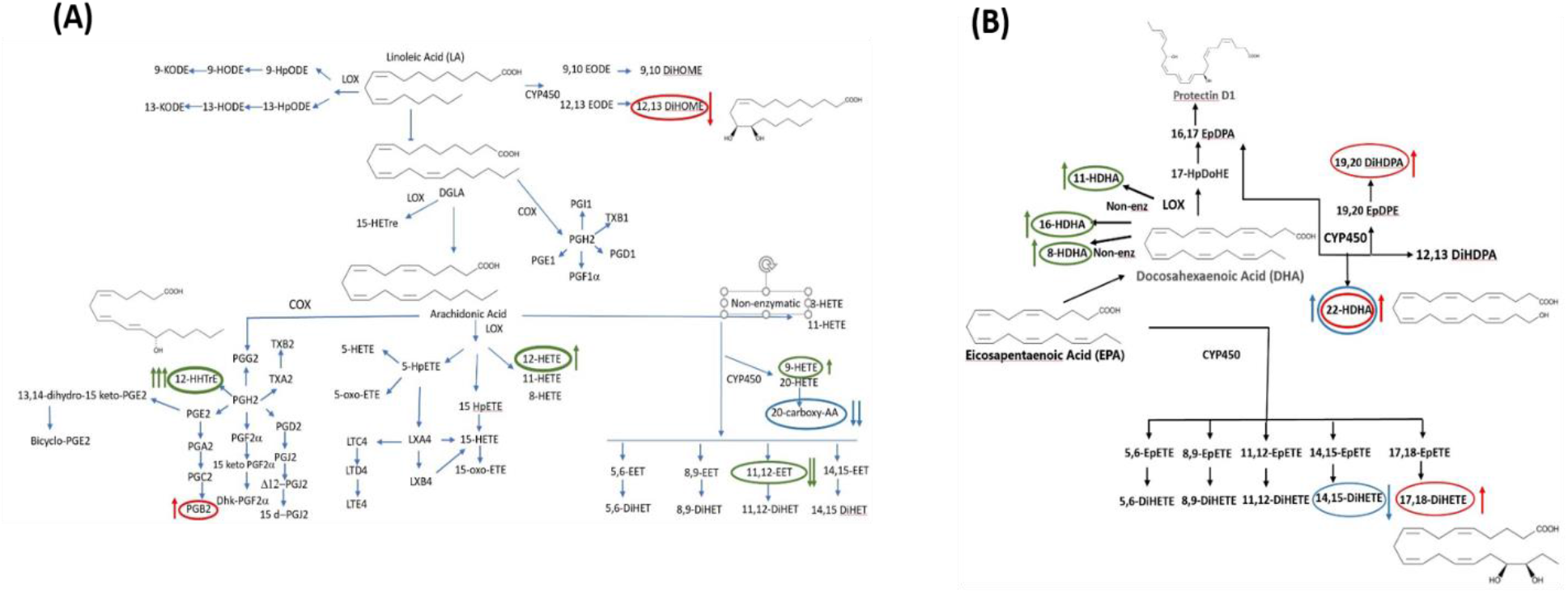
Radiation induced eicosanoid changes in l6weeks old ApoE^-/-^ mice. **(A)** illustrates eicosanoid pathways contributing to the heart base vs no RT effect (green circles), contrasted with apex vs no RT effect (red circles) all downstream from linoleic acid or arachidonic acid. (B) illustrates eicosanoid pathways contributing to the heart base vs no RT effect (green circles), contrasted with apex vs no RT effect (red circles) and base vs apex effects (blue circles) all downstream from EPA or DHA.

In this study, using ApoE ^-/-^ mice on HFD subjected to 16Gy irradiation of the whole heart or partial irradiation (base or apex) atherosclerotic lesions formation and inflammatory changes in response to RT were assessed. We found that the base of the heart is significantly more prone to the development of atherosclerotic lesions post-RT. In addition, there was an increase of inflammatory cells in the irradiated areas of the heart, indicative of an endothelial inflammation mediated response in the development of RT-related CVD.

## Methods

### Experimental design

Male ApoE^-/-^ mice on a C57BL/6j background were purchased from Jackson Laboratories (Bar Harbor, ME). The mice were housed at the Research Animal Resource Center housing facility of Memorial Sloan Kettering Cancer Center, a facility approved by the American Association for Assessment and Accreditation of Laboratory Animal Care and maintained in accordance with the regulations and standards of the United States Department of Agriculture and the Department of Health and Human Services, NIH. Protocols for conducting animal experiments were approved by the MSKCC Institutional Animal Care and Use Committee (IACUC).

The animals were divided in two experimental groups: (a) Eight-week old mice (5 mice each in the no RT/control, whole heart RT, basal or apical RT group) were placed on a high-fat diet [0.2% total cholesterol + Total fat (21% by weight; 42% kcal from fat) + >60% of total fatty acids + High sucrose (34% by weight)= HFD] (Envigo, IN, USA) and irradiated as mentioned above at 9 weeks of age. Eight weeks post irradiation, the mice were euthanized and hearts and aortas were harvested for subsequent analysis. (b) Seven-week old mice (5 mice each in the no RT/control, whole heart RT, basal or apical RT group) were placed on HFD and irradiated as mentioned above at 16 weeks of age. Eight weeks post irradiation, the mice were euthanized and the hearts and aortas were harvested for subsequent study.

### Cardiac irradiation

Mice were anaesthetized with Isoflurane and irradiated using a 225C (Precision X-Ray, North Branford CT), 10×10mm collimator, at 225kV, 13mA, at a dose rate of 3.5Gy/minute. A cone beam computed tomography (CBCT) scout was performed on each mouse followed by a CBCT scan. After the CBCT scan, the target of irradiation was selected using the PilotXRad Application. Thus, precise regions of cardiac irradiation – whole, base or apex of the heart were selected and irradiated with one total dose of 16Gy.

### Validation of the irradiated areas of the heart

One hour after irradiation, mice were euthanized using 100% carbon dioxide at 5 PSI for 3 minutes and left undisturbed for an additional 15 minutes. Hearts were perfused with PBS using a perfusion chamber and then harvested. Hearts were fixed in 10% formalin overnight at 4°C. The hearts were embedded in paraffin and sections were cut and immuno-stained for γ-H2AX foci. Radiation-induced double strand breaks were identified by the generation of γ-H2AX foci and served as a biological marker for the irradiated field. The anatomical distribution of the foci in the heart was related to the field size and positioning to provide information on the precision of the field exposed.

### Heart tissues immunohistochemistry (IHC)

The hearts were harvested 8 weeks post-RT, fixed in 10% neutral buffered formalin (NBF) for 48 hours and cut transversally at the mid-horizontal plane. Both halves were placed into one tissue cassette with the planar cut-surface facing downwards. The tissue was processed in alcohol and xylene and embedded in paraffin. The tissue blocks were sectioned at 4 μm thickness at up to 20 levels per heart with an interlevel width of 144 μm, in order to sample the entire ventricular myocardium. Each level was stained with hematoxylin and eosin (H&E).

At level 8, six additional unstained serial sections were performed to allow further characterization with special stains or IHC. IHC for CD3^+^ cells (Abcam ab135372, 1: 250 following heat-induced epitope retrieval [HIER] in a pH 6.0 buffer), CD45^+^ cells (BD Pharmingen 550539, 1:250, HIER pH 6.0), and CD31^+^ (Dianova DIA-310, 1:250, HIER pH 6.0) was performed on sections of level 8 of each heart. All stains were performed in a Leica Bond RX automated stainer using the Bond Polymer Refine detection system (Leica Biosystem DS9800). The chromogen used was 3,3 diaminobenzidine tetrachloride (DAB) and sections were counterstained with hematoxylin. An average of 20 HE slides per heart was analyzed. All H&E slides were evaluated, and coronary atherosclerotic lesions were counted separately for the base and apex at each level for each heart, and additional lesions in the myocardium were scored semi-quantitively and recorded by a board-certified veterinary pathologist (S. Kitz).

### Digital Image Analysis and Cell Quantification

The CD3 and CD45 expression on IHC of inflammatory cell infiltrates were evaluated quantitatively by automated image analysis. Microvascular density (MVD) was quantitatively evaluated by measurement of CD31 ^+^ cells. Both quantifications were done on sections from IHC performed on level 8 for every heart. Whole-slide digital images were generated on a scanner (Pannoramic 250 Flash III, 3DHistech, 20x/0.8NA objective, Budapest, Hungary) at a resolution of 0.2431 μm per pixel. Image analysis was performed utilizing QuPath software [25]. The region of interests (ROIs) were manually defined as the entire ventricular myocardium section from the base and apex (analyzed separately) on each image, excluding valves, auricles and epicardial connective tissue. The positive cell detection algorithm was used to detect CD3^+^ and CD45^+^ cells and categorize them as positive, based on the optical density (OD) of DAB in the cytoplasm. ROI selection, positive cell detection algorithm optimization, OD threshold determination, and validation of the results were performed by a board-certified veterinary pathologist (S. Kitz). For CD3^+^ and CD45^+^ the number of positive cells per mm^2^ were counted separately for the base and apex of each image. For the quantification of the microvascular density, the number of vessels per mm^2^ was counted separately for the base and apex of each image, and percentage of CD31^+^ pixels was measured and calculated for base and apex of each image using the pixel classifier tool.

### Aorta harvesting and staining

The extent of atherosclerosis in the aorta was assessed by *en-face* analysis of the whole aorta [26–29]. Entire aortas (thoracic and abdominal) were harvested from all mice 8 weeks following RT. Aortas were fixed in 10% NBF at 4°C following which they were stored in PBS. For *en-face* staining, the whole aorta was placed onto a tray containing smoothened black wax and covered with PBS. The aorta was gently cleaned under a dissection microscope to remove all loose perivascular adipose tissue around the aorta from the thorax and abdomen. The aorta was the pinned on the wax board with 0.15mm minutien pins to open an end of the common iliac artery. The aorta was then cut longitudinally using spring scissors to allow complete splitting open of the aorta and then pinned down to allow them to stay flattened.

Prior to staining the aorta, 1% stock solution of Oil Red O (ORO) (Sigma Aldrich 00625-100g) was prepared in isopropyl alcohol. A 2:3 working solution was prepared by diluting the stock in deionized water and the solution was filtered before use. Each aorta was rinsed in 70% isopropyl alcohol followed by ORO staining and then rinsed again in 70% isopropyl alcohol. The aortas were then placed in PBS and images taken using a Lumar stereoscope attached to a camera.

### Digital image analysis

The digital images of the entire Oil Red O stained aortas were evaluated using image analysis software Fiji [30]. The aortic atherosclerotic lesions were expressed as a percentage of the area of lesions (red), relative to the surface area of the entire aorta. All continuous variables were expressed as mean ± standard error of mean.

### Biochemical Eicosanoid Measurements

~100 eicosanoids were measured 8 weeks post-RT from the serum of 16 weeks old ApoE ^-/-^ mice. Serum from each mouse from the different treatment groups of mice (non-irradiated, whole heart RT, basal or apical heart RT) was used to measure eicosanoids by liquid chromatography mass spectrometry techniques using the Sciex 6500+ system^®^. Quality control samples were made, after a portion of all samples were pooled and were assessed every 10 samples for calibration of signal intensities.

### 2D-Ultrasound

Echocardiography were performed on 16 weeks old mice that receive radiation to whole, base or apex of the heart, one total dose of 16Gy at 4 and 8 weeks post-RT. Images were acquired using the Vevo 2100 Imaging System. Acquisitions were made using the Brightness (B) mode to acquire two dimensional images in the Parasternal Long Axis as well as short axis View. Cardiac parameters such as ejection fraction were calculated using the imaging software.

### Statistics

Data are expressed as mean ± SEM. Statistical analysis were performed using One-way ANOVA and post hoc Multiple Tukey’s comparison test (Prism 9.0.0. GraphPad). Significance was set at p < 0.05.

## Results

### Validation of irradiation specific regions of the heart

It has been previously shown that the formation of atherosclerotic lesions in ApoE^-/-^ mice is accelerated by cardiac irradiation. It has also been demonstrated that atherosclerotic lesions in these mice increase significantly in a radiation-dose dependent manner [16]. Cardiac irradiation of ApoE^-/-^ mice with 16Gy led to a significant increase in the number of coronary atherosclerotic lesions in the hearts at 20 weeks old mice post-RT as compared to unirradiated control mice or mice receiving 2Gy or 8Gy cardiac irradiation [16]. Thus, for the studies described here we used 16Gy to further elaborate the relationship between the region of cardiac irradiation and development of CVD. The previous studies have been performed using standardized chow [16, 31]. In the studies described here we used ApoE^-/-^ mice on HFD to understand the interplay of RT, site of irradiation, age and diet on the development of RT-induced CVD.

In order to accurately irradiate specific regions of the heart – whole heart, base or apex of the heart the small animal 225C Precision X-ray irradiator and 10×10mm collimator were used. This irradiator has a Computed Tomography scanning feature which allows to locate specific parts of the heart and to deliver 16Gy to the selected field of irradiation with a high degree of precission. The different irradiated fields were confirmed, by stainning for γ-H2AX foci, as a biological marker for RT-induced double strand breaks. As shown in Fig. 1, the hearts of mice receiving whole heart irradiation, showed γ-H2AX foci in both the basal and apical regions of the heart. In mice receiving basal or apical cardiac irradiation only, γ-H2AX foci were observed only in the base or the apex regions of the heart respectively. Mice that served as controls and therefore received no irradiation, their hearts and lungs from all experimental groups showed no presence of γ-H2AX foci, thus confirming the accuracy of the radiation treatment plan using the 225XR Precision X-ray irradiator.

### RT treatment toxicity

Mice involved in this study were monitored for toxicity of the treatments by measuring their weight twice a week up to 8 weeks following RT. The weights of 9 week-old control mice on HFD that received no irradiation were not significantly different from the age-matched mice that received basal cardiac irradiation for up to 8 weeks post-RT. Apical cardiac irradiation led to a significant reduction in body weight beginning at 5 weeks post-RT as compared to non-irradiated controls. Whole heart irradiation resulted in a significant reduction in body weight at 8 weeks post-RT. The heart weights in the different experimental groups were not significantly different at 8 weeks post-RT (data not shown). No significant differences were observed in the body weight or heart weight at 8 weeks post-RT between experimental groups of no irradiation, whole heart, apical or basal irradiation in ApoE ^-/-^ mice that received irradiation at 16weeks of age (data not shown).

### Cardiac hypertrophy

Ratios of heart weight to tibia length and heart weight to body weight were measured to assess whether irradiation and HFD led to cardiac hypertrophy in ApoE^-/-^ mice. No significant differences were observed in either ratios in both the 9 weeks or 16 weeks old mice that received no RT, whole heart, apical or basal RT (data not shown). These results indicate that the differential cardiac irradiation did not induce cardiac hypertrophy.

### Coronary atherosclerosis

In the first set of experiments 8 weeks ApoE^-/-^ mice on HFD received whole, basal or apical cardiac RT at 9 weeks of age. We observed a significant increase in the number of basal atherosclerotic lesions in the myocardial and sub-endocardial vasculature of the mice receiving whole heart RT (Fig.2A right panel). The number of lesions seen in the basal myocardial vasculature post basal RT (7.8 ± 2.49) were not significantly different than those observed post-whole heart RT (12.2 ± 3.29). There was no significant difference in the number of lesions in the myocardial vasculature at the base post-apical RT (2 ± 0.948) as compared to control (0). The number of atherosclerotic lesions in the coronary arteries at the base of the heart in control mice (10.8 ± 2.53), whole heart-RT (11 ± 2.88), apical-RT (5.2 ± 1.77) or basal-RT (13.2 ± 3.96), were not statistically significantly different from each other (Fig. 2A left panel).

The number of lesions seen in the apical coronary arteries post-whole heart RT (9.6 ± 3.29) were significantly higher than in the unirradiated controls (0.8 ± 0.58) or apical irradiated group (1.8 ± 1.31) (Fig. 2B left panel). The response of the coronary arteries in the apex of the heart to whole heart or basal irradiation (5 ± 1.22) led to comparable number of atherosclerotic lesions. Although there was a trend towards an increase number of atherosclerotic lesions in the myocardial and sub-endocardial vasculature in the apex post-whole heart RT (10.8 ± 5.43) or post-apical RT (8 ± 2.62) as compared to control or post-basal RT (0.4 ± 0.24), these differences were not significantly different (Fig. 2B right panel).

Next, we assessed the combined effect of age as well as RT on the development of atherosclerotic lesions. ApoE^-/-^ mice, 7 weeks old, were kept on HFD and received whole, basal or apical irradiation in the heart at 16 weeks of age. Hearts were harvested 8 weeks post-RT and the formation of atherosclerotic lesions was analyzed as previously described. The basal lesions formed in the coronary arteries of control mice (9 ± 2.70) were lower but not statistically significantly different from those formed post-whole heart RT (17.6 ± 2.11). (Fig. 3A left panel). A significant increase in the atherosclerotic lesions of the basal coronary arteries was observed after irradiation of the base of the heart (29.33 ± 5.48) as compared to non-irradiated controls. In response to apical cardiac irradiation, the number of atherosclerotic lesions in the coronary arteries at the base of the heart (11 ± 3.06) remained comparable to those seen in the controls (Fig. 3A left panel). The atherosclerotic lesions in the myocardial and sub-endocardial region of the base, after basal-RT (6.66 ± 2.07) were significantly greater as compared to control mice (0.2 ± 0.2) (Fig. 3A right panel). Thus, mice receiving basal-RT at 16 weeks of age on HFD demonstrated the highest number of atherosclerotic lesions in the base of the heart, further demonstrating the sensitivity of this area of the heart to irradiation.

The coronary arteries in the apical region of the heart, did not respond differently to control, whole heart or partial cardiac irradiation as seen by comparable number of atherosclerotic lesions formed in these groups (Fig. 3B left panel). Whole heart (7 ± 1.51) and apical (6.8 ± 1.06) irradiation led to a significant increase in atherosclerotic lesions formed in myocardial and subendocardial vasculature at the apex as compared to unirradiated controls (0.6 ± 0.4) (Fig. 3B right panel).

### Inflammatory changes in irradiated hearts

Evidence of inflammation was assessed in the age-matched differentially irradiated ApoE ^-/-^ mice kept on HFD by evaluating the infiltrating CD45^+^ and CD3^+^ cells using IHC. At 8 weeks post-whole heart RT of 9 weeks old mice, there was a significant increase in the number of CD45^+^ cells at the base (102.36 ± 11.36 per mm^2^) (Fig. 4A) and apex (98.66 ± 17.28 per mm^2^) (Fig. 4A) of the hearts as compared to non-irradiated controls (21.56 ± 3.48 per mm^2^ in the base and 19.60 ± 17.28 per mm^2^ in the apex). Basal-RT led to a significant increase in CD45^+^ cells at the base of the hearts (171.39 ± 19.12 per mm^2^) as compared to unirradiated controls, whole heart RT or, apex RT (Fig. 4A). Conversely, apical irradiation led to a significant increase CD45^+^ positive cells in the apex of the hearts (145.65 ± 15.05 per mm^2^) when compared with CD45^+^ cells present in the apex of the hearts of the animals from unirradiated controls, whole heart irradiated or basal-RT.

The infiltration of the CD3^+^ T cells showed a similar pattern to the one observed for the CD45^+^ cells infiltration in the different irradiated parts of the heart. The number of CD3^+^ T cells also significantly increased in the base (11.37 ± 2.17 per mm^2^) and apex (11.10 ± 2.45 per mm^2^) regions of mouse hearts 8 weeks post-whole heart RT as compared to non-irradiated controls (Fig. 4B). Basal-RT led to a significant increase in CD3^+^ T cells only in the basal region of the hearts (11.05 ± 1.64 per mm^2^) compared to CD3^+^ T cells observed in the base of the hearts of non-RT or apical-RT mice. Mice that received apical cardiac irradiation demonstrated a significantly greater number of CD3^+^ T cells in the apex (8.95 ± 1.29 per mm^2^) as compared to those present in the apex of control or basal irradiated mice (Fig. 4B). Interestingly, the number of CD3^+^ T cells observed in the base of the hearts of mice after whole heart or basal irradiation was comparable. Similarly, the CD3^+^ T cell numbers observed in the apex of the hearts of mice after whole heart or apical irradiation were not significantly different from each other (Fig. 4B).

Figures 4C and 4D show the infiltration of CD45^+^ and CD3^+^ T cells in ApoE ^-/-^ mice at 16 weeks of age on HFD who received whole, basal or apical RT. Hearts were harvested 8 weeks post-RT to analyze the infiltration of these cells. Similar to the finding in the 9 weeks old mice, the expression of CD45^+^ and CD3^+^ T was restricted locally to the region of irradiation. Whole heart RT led to a significant increase in the CD45^+^cells in the base (382.17 ± 64.59 per mm^2^) and the apex (300.15 ± 62.12 per mm^2^) as compared to non-RT controls (117.54 ± 32.51 per mm^2^ in the base and 67.67 ± 8.24 per mm^2^ in the apex) (Fig. 4C). Basal irradiation led to an increase in the number of CD45^+^ cells in the base (594.95 ± 33.95 per mm^2^) which was greater as compared to control, whole heart or apical RT (148.13 ± 23.08 per mm^2^). The CD45^+^ cells post-apical RT remained comparable to unirradiated controls (Fig. 4C).

The number of CD3^+^ T cells in the base was significantly increased in mice that received whole heart (43.50 ± 8.05 per mm^2^) or basal-RT (42.13 ± 1.64 per mm^2^) as compared to unirradiated controls (7.14 ± 2.41 per mm^2^) (Fig. 4D). Whole heart (42.11 ± 9.37 per mm^2^) and apical-RT (33.11 ± 4.63 per mm^2^) led to a significant increase in the CD3^+^ T cells in the cardiac apex.

### Basal Heart Irradiation Induces Pro-Atherogenic Eicosanoid Mediator Effects

Our data on RT of the base of the heart of 16 weeks old ApoE^-/-^ mice shows significantly more pro-atherogenic changes in the COX, CYP450 and LOX pathways (Fig.5 and Table 1). As shown in Table 1, radiation to the heart base vs no RT or apex, results in more eicosanoid species with a higher fold changes than for apex RT vs no RT, and consistent with the pronounced effect of base heart radiation on atherosclerosis development. One of the major treatment targets of atherosclerotic and ischemic heart disease is the COX pathway, which modulates major pathophysiological features of these diseases, including platelet aggregation, vessel wall tension, and inflammatory processes in atherosclerotic lesions [32]. The most highly elevated of the COX metabolites (pro-inflammatory) was seen for base radiation vs no RT, 12(S)-HHTrE (8-fold, Table 1), one of the primary arachidonic acid (AA) metabolites of human platelets, accounting for approximately one third of the AA metabolites produced by stimulated platelets may thus be an important local modulator of platelet plug formation [33]. It’s biosynthesized by thromboxane (TXA2) synthase from prostaglandin H2 (PGH2). CYP450 contribution to the base irradiation effect of atherosclerosis may include the 3-fold decrease of 11,12 EET (11,12-EpETrE), as well as the 1.6 fold increase in 9-HETE, and the 35% decrease in 14,15 DiHETE. 11,12 EET has vasodilatory effects in intestinal microcirculation [34] and dilates coronary microvessels in canine and porcine models [35], and inhibits vascular inflammation, distinct from its vasodilatory effects by inhibiting NF-kB and inhibitor of k B kinase in murine model [36]. Levels of HETEs were significantly elevated in plaques retrieved from symptomatic patients compared with those retrieved from asymptomatic patients, and the most abundant was found to be 9-HETE, which accounted for 21.4 ± 1.9% of the total HETEs [37]. EPA derived 14,15DiHETE (35% decreased in base vs apex RT) inhibits human platelet aggregation, but with much less potency than parent EpETE [38]. Lipoxygenase contribution to the base irradiation effect of atherosclerosis may include 12-HETE (1.6 fold elevated), which has been seen to increase monocyte adhesion to human endothelial cells leading to aortic fatty streak formation [39, 40]. Oxidative stress is clearly evidenced by the chronic elevation in base heart RT vs sham treated of non-enzymatic oxidation products of DHA seen, 8-HDHA,11-HDHA and 16-HDHA, all approximately 50% elevated (Table 1).

**Table 1.** Fold changes of eicosanoid species and pathways, elicted by positional heart irradiation (apex vs heart base). Pathways are further described, by matched color coding of species shown to circles in Fig. 5A and Fig. 5B. Abbreviations: LA-linoleic acid 18:2 (n-6); EPA - Eicosapentaenoic acid (20:5(n-3)), DHA-Docosahexaenoic acid 22:6(n-3), AA-arachidonic acid (20:4, (n-6)); CYP450-Cytochrome p450; COX-Cyclooxygenase; LOX-arachidonate-(5, 12 or 15) lipoxygenase non-enz nonenzymatic (autooxidation).

| Species | Formula | Pathway | Ratio | Fold Change | t-test |
| --- | --- | --- | --- | --- | --- |
| 12,13-DiHOME | C<sub>18</sub>H<sub>34</sub>O<sub>4</sub> | LA->CYP450 | Apex/no RT | 1.538073 | 0.0038 |
| 17,18-DiHETE | C<sub>20</sub>H<sub>32</sub>O<sub>4</sub> | EPA-><br>CYP450 | Apex/no RT | 1.3487 | 0.016 |
| 19(20)-DiHDPA<br>19(20)-DiHDPE | C<sub>22</sub>H<sub>34</sub>O<sub>4</sub> | DHA-><br>CYP450 | Apex/no RT | 1.25903 | 0.023 |
| Prostaglandin B2 | C<sub>20</sub>H<sub>30</sub>O<sub>4</sub> | AA-> COX | Apex/no RT | 1.555194 | 0.027 |
| 22-HDHA | C<sub>22</sub>H<sub>32</sub>O<sub>3</sub> | DHA-><br>CYP450 | Apex/no RT | 0.777624 | 0.049 |
| 12(S)-HHTrE<br>12(S)-HHT | C<sub>17</sub>H<sub>28</sub>O<sub>3</sub> | AA->COX | Base/no RT | 8.783465 | 0.006 |
| 9-HETE | C<sub>20</sub>H<sub>32</sub>O<sub>3</sub> | AA->CYP450 | Base/no RT | 1.610849 | 0.026 |
| 16-HDHA<br>16-HDoHE | C<sub>22</sub>H<sub>32</sub>O<sub>3</sub> | DHA->non-enz | Base/no RT | 1.56499 | 0.027 |
| 12-HETE | C<sub>20</sub>H<sub>32</sub>O<sub>3</sub> | AA->LOX | Base/no RT | 1.638786 | 0.027 |
| 8-HDHA<br>8-HDoHE | C<sub>22</sub>H<sub>32</sub>O<sub>3</sub> | DHA->non-enz | Base/no RT | 1.635577 | 0.035 |
| 11-HDHA<br>11-HDoHE | C<sub>22</sub>H<sub>32</sub>O<sub>3</sub> | DHA->non-enz | Base/no RT | 1.466157 | 0.05 |
| 11,12-EET | C<sub>20</sub>H<sub>32</sub>O<sub>3</sub> | AA->CYP450 | Base/no RT | 0.383295 | 0.05 |
| 22-HDHA | C<sub>22</sub>H<sub>32</sub>O<sub>3</sub> | DHA->non-enz | Base/Apex | 1.464036 | 0.05 |
| 20-carboxy<br>Arachidonic Acid | C<sub>20</sub>H<sub>30</sub>O<sub>4</sub> | AA->CYP450 | Base/Apex | 0.266334 | 0.05 |
| 14(15)-DiHETE | C<sub>20</sub>H<sub>32</sub>O<sub>4</sub> | EPA->CYP450 | Base/Apex | 0.658104 | 0.05 |

Apex irradiation eicosanoid mediators are not clearly atherogenic as those from base irradiation. 12,13-DiHOME (50% elevated vs no RT) is known to cause mitochondrial dysfunction [41], 17,18-DiHETE (35% elevated vs no RT), and 19,20-DiHDPE (25% elevated vs no RT) inhibits human platelet aggregation [38]. Prostaglandin B2 (55% elevated vs no RT) is known to mediate mesenteric vascular dose-dependent vasodilatory and vasoconstrictory effects in animal models [42]. 22 HDHA (22 hydroxy DHA) is simply a non-enzymatic oxidation production of DHA, and is elevated more with base heart irradiation vs apex (46% increased in base vs apex irradiation, 22% decreased in apex vs no RT).

### Microvascular changes post differential cardiac irradiation

We next assessed changes in the microvascular density by measuring CD31 expression using IHC staining of cardiac sections harvested 8 weeks post whole heart, apical or basal-RT as compared to unirradiated control. In the ApoE ^-/-^ mice irradiated at 9 weeks of age, whole heart-RT led to a significant increase in CD31^+^ cells per mm^2^ of cardiac area in the base (183.13 ± 20.87 per mm^2^) as well as the apex (168.08 ± 33.85 per mm^2^) (Fig. 6A). Basal RT had no impact on the CD31^+^ cells in the apex (25.82 ± 4.65 per mm^2^) as compared to unirradiated controls (21.87 ± 10.49 per mm^2^) but showed a significant increase in the CD31 ^+^ cells in the base (166.29 ± 10.69 per mm^2^) as compared to controls (39.65 ± 18.69 per mm^2^) (Fig. 6A). Similarly, apical RT significantly increased the CD31 ^+^ cells in the apex (180.84 ± 20.79 per mm^2^) as compared to unirradiated region apical CD31 ^+^ cells (21.87 ± 10.49 per mm^2^).

**Figure 6.**
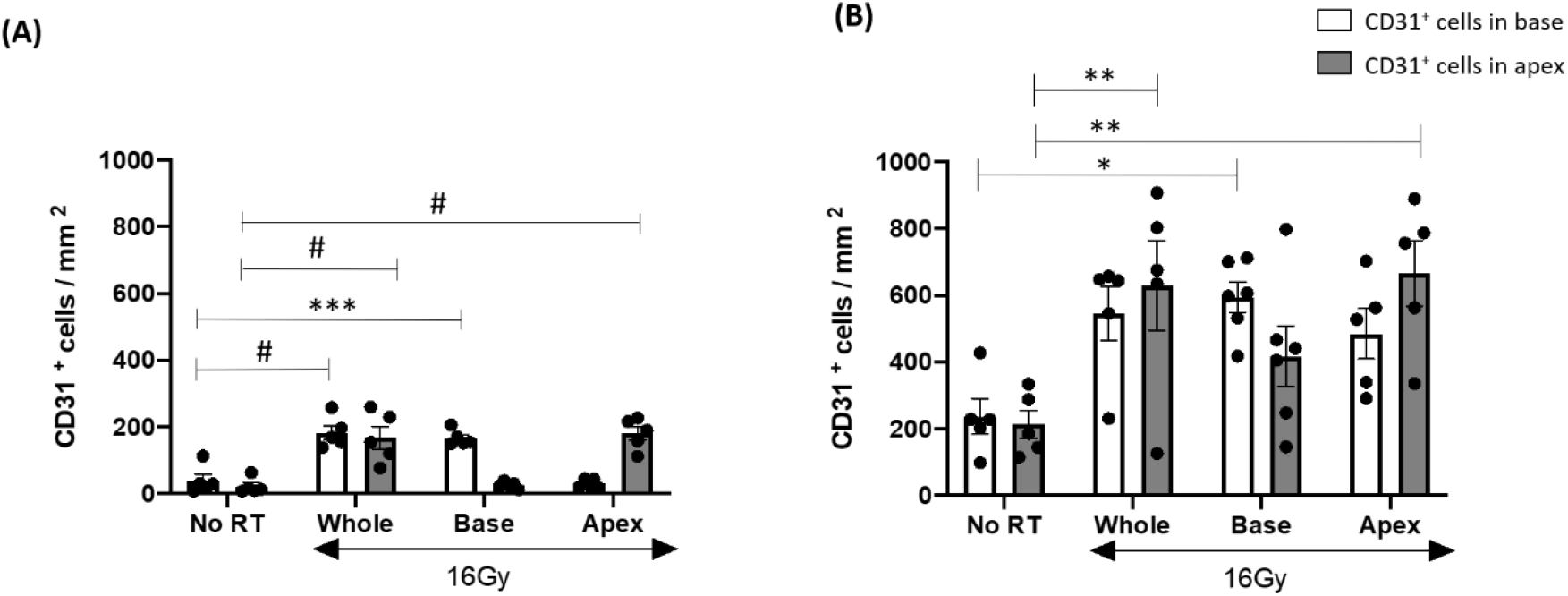
CD31 changes observed post differential cardiac irradiation of ApoE^-/-^ mice. (A) CD31^+^ cells observed in the base and apex of 9 weeks old mice after whole heart, basal, apical or no irradiation. (B) CD31^+^ cells observed in the base and apex of 16 weeks old mice at 8 weeks follow-up after whole heart, basal, apical or no irradiation. Bars represent mean ± SEM. * p<0.05, **p<0.01, ***p<0.001, # p<0.0001. n=5 per group, each dot represents number of CD31^+^ cells from the heart of one mouse.

At 8 weeks post-RT of 16 weeks old mice, there was a significant increase in the number of CD31 ^+^ cells in the basal region only of hearts from mice that received basal irradiation (594.71 ± 33.8 per mm^2^) (Fig. 6B). These mice already demonstrated a greater number of CD31 ^+^ cells even in the unirradiated controls (236.84 ± 53.37 per mm^2^ in the base and 213.69 ± 41.96 per mm^2^ in the apex) (Fig. 6B), thus indicating that differential cardiac irradiation may have further serious implications in the setting of HFD and age. Whole heart (629.51 ± 134.59 per mm^2^) and apical irradiation (666.00 ± 98.14 per mm^2^) led to a significant increase in the CD31^+^ cells observed in the apex of the mice (Fig. 6B).

### Aortic Atherosclerotic lesions

The thoracic and abdominal aortas from the two sets of experiments described above were used to assess the effect of age and differential cardiac irradiation on the development of atherosclerotic lesions in the aorta while the mice were on HFD (Fig. 7). A ratio of the area of atherosclerotic lesions relative to the surface area of the entire aorta was compared in each cardiac irradiation group and also each experimental group. Male ApoE ^-/-^ mice at 8 weeks of age were kept on HFD and received whole, basal or apical RT in the heart at 9 weeks of age. Follow up 8 weeks post-RT, showed no significant difference between aortic lesions in the whole heart, apical or basal irradiation groups as compared to unirradiated control.

**Figure 7.**
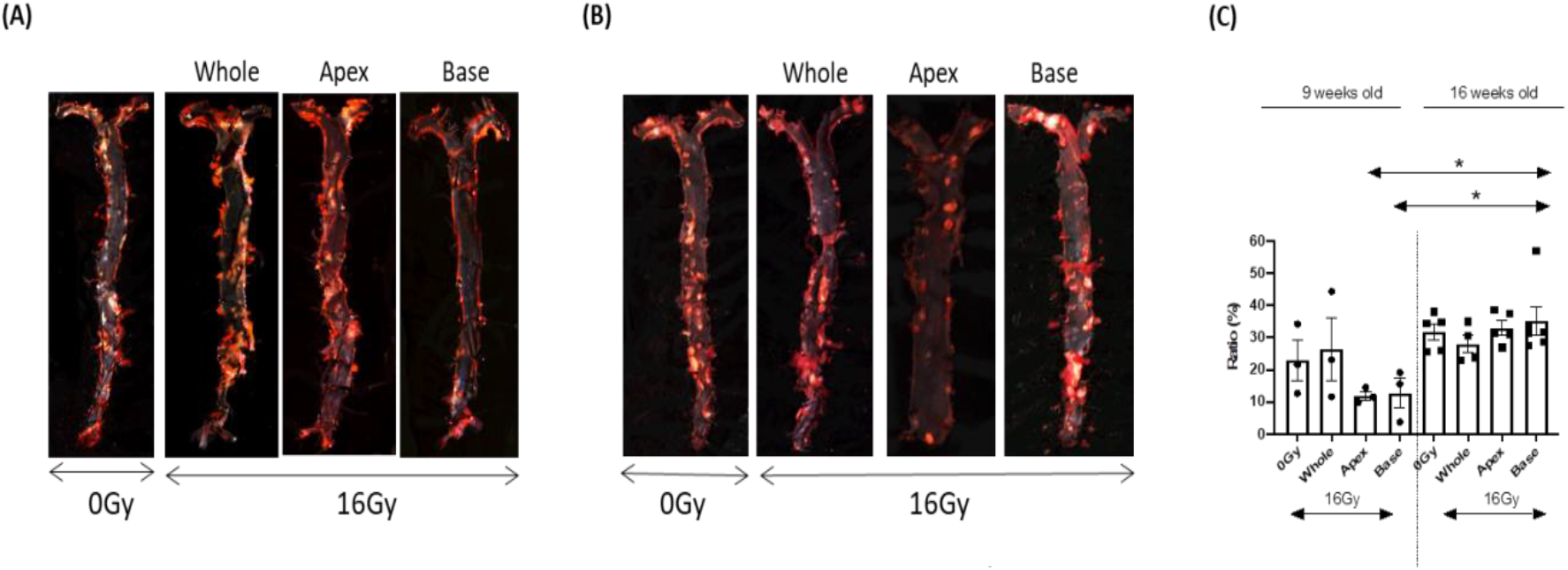
Aortic atherosclerotic lesions 8 weeks post cardiac irradiation of ApoE^-/-^ mice. **(A)** Aortic lesions observed in 9 weeks old mice after whole heart, basal, apical or no irradiation (B) Aortic lesions observed in 16 weeks old mice after whole heart, basal, apical or no irradiation (C) Quantification representing the ratio of the area of atherosclerotic lesions relative to the surface area of the entire aorta. Each dot represents the aorta from one mouse. Bars represent mean + SEM. * p<0.05.

Male ApoE^-/-^ mice, 7 weeks old, on HFD received whole, basal or apical RT at 16 weeks of age. Similar to previous results obtained at 8 weeks post-RT, in 9 week old mice, there was no significant difference between whole heart, apical or basal cardiac irradiation and unirradiated controls. The number of aortic lesions formed in the control, whole or apical cardiac irradiation were comparable in the younger (9 weeks old) and older (16weeks old mice). However, basal irradiation led to a significant increase of aortic atherosclerotic lesions in 16 weeks old mice compared to those formed by basal cardiac irradiation in 9 weeks old mice (39.87% vs 12.40%) indicating that age is an important component in the RT-induced acceleration of aortic atherosclerotic lesions in the heart.

### Cardiac function

Ejection fraction was used as an assessment of global cardiac function measured by cardiac ultrasound at 4 or 8 weeks post-RT on ApoE ^-/-^ mice on HFD from 7 weeks of age and irradiated at 16 weeks with 16Gy. There were no significant changes in cardiac ejection fraction among the experimental groups. Other parameters such as stroke volume, cardiac output, endocardial fractional area change or fractional shortening between the experimental groups of mice that received no, whole heart, basal or apical irradiation remained unchanged in spite of RT-induced acceleration of atherosclerotic lesion formation (data not shown).

## Discussion

Modern day RT involves highly specialized radiation techniques that are successful in excluding irradiation of normal tissues, however partial irradiation and systemic RT effects received are still a significant clinical problem [43]. To the best of our knowledge, this is the first experimental study mimicking clinical RT for cancer patients wherein only a specific region of the heart is included in the radiation field to assess the sensitivity of different areas of the heart. Our findings support the hypothesis that distinct regions of the heart respond differentially to radiation exposure in terms of the development of vascular atherosclerotic lesions. We have previously shown that the formation of atherosclerotic lesions is increased in ApoE ^-/-^ mice on a regular chow diet by cardiac radiation [31] and studies by others have demonstrated that these effects are radiation-dose dependent [16].

In this study we demonstrate that in 9 weeks old ApoE^-/-^ mice on HFD: (1) Basal and whole cardiac RT led to comparable number of atherosclerotic lesions in the myocardial and sub-endocardial vasculature, whereas apical irradiation had no impact on the formation of these lesions; (2) The coronary arteries of the apex developed a significantly greater number of atherosclerotic lesions after whole heart irradiation as compared to unirradiated controls. The coronary apical lesions after whole heart or basal RT were however not significantly different from each other.

Different findings were observed in the in 16 weeks old ApoE ^-/-^ mice on HFD: (3) Basal irradiation led to a significant increase in the number of basal atherosclerotic lesions in both the coronary arteries and the myocardial and subendocardial vasculature; (4) Significant atherosclerotic lesions were observed in the apical myocardium following whole and apical cardiac RT versus unirradiated controls; (5) In the 9 weeks old ApoE^-/-^ mice the increased infiltration of the inflammatory cells correlated with the irradiated area of the heart. The number of infiltrating inflammatory cells at baseline was increased for both CD45^+^ and CD3^+^ T cells in the older mice, which might be explained by a chronic inflammatory process that evolved with the increased age of these ApoE^-/-^ mice while on HFD. In addition, there was a significant increase in the number of CD45^+^ cells infiltrating in the base of the heart post-base RT but not in the apex in response to apex RT, indicating that in these mice indeed the base of the heart is more susceptible to inflammation than the apex; (6) Pro-inflammatory and pro-atherogenic eicosanoid pathways were stimulated post basal cardiac irradiation as seen by the up-regulation of (±)11(12)-EET and down-regulation of 12(S)-HHTrE, (±)9-HETE, (±)16-HDHA, (±)12-HETE, (±)8-HDHA and (±)11-HDHA. (7) No differences were observed in the development of atherosclerotic lesions in the aorta between the different areas of the heart irradiated as compared to unirradiated controls. However, in the older (16 weeks old) mice the number of aortic atherosclerotic lesions post basal-RT were significantly higher than those observed in the 9 weeks old mice after basal RT.

Clinical studies have shown that patients receiving thoracic RT have an increased risk of coronary artery disease, valvular heart disease, congestive heart failure, pericardial disease and sudden death [17] and experimental studies have shown pre-lesion changes in the descending thoracic aorta of wild-type mice after total body irradiation (TBI, single acute exposure of 5Gy) [44]. It has been suggested that changes observed in the thoracic aorta following TBI could result not only from direct effects to the thoracic aorta but also from effects from various adjacent organs/tissues [44].

Studies using hypercholesterolemia (ApoE^-/-^) and local cardiac irradiation have shown that accelerated development of coronary atherosclerosis is a combination of an inflammatory response, microvascular and endocardial injury [16]. In these studies mice showed an increased influx of CD45^+^ and CD3^+^ T cells in the myocardium 40 weeks after 8 and 16Gy cardiac irradiation [16]. In our studies, the expression of both CD45^+^ and CD3^+^ T cells was associated with the location of cardiac irradiation. Whole heart irradiation led to increased CD45^+^ cells in both the base and apex of the heart, whereas partial irradiation in the apex or base led to significant increase in CD45^+^cells only in the apex or base respectively. Therefore, we speculate that the expression of an inflammatory response, such as infiltration of CD45^+^ and CD3^+^ T cells, might contribute to the progression of RT-induced CVD. This inflammatory process was confirmed also by the regulation of eicosanoids found in the serum of 16 weeks old ApoE ^-/-^ mice exposed to RT. It is of interest that base irradiation results in the detection of proatherogenic eicosanoids mediators in serum, or the decease in eicosanoid mediators that inhibit platelet aggregation, inflammation or vasodilatation (see Table 1). In support of a role of eicosanoid mediators for base or whole heart atherogenic irradiation effects, apex irradiation eicosanoid mediators are not clearly atherogenic, in contrast to eicosanoid mediators detected in serum after base heart irradiation.

As the increase in the acute and/or chronic inflammatory responses were detected in the cardiac base when mice received whole or partial basal irradiation, it is possible that the similar increase in atherosclerotic lesions after whole heart or basal irradiation is associated with the increased CD45^+^ and CD3^+^ T cells infiltration. Further studies are necessary to fully establish the inflammatory effects of RT in the partial exposure of the heart as it pertains to both eicosanoids pathways and cytokines/chemokines secretion [45].

Older mice on HFD, which received differential cardiac irradiation showed an increase in CD45^+^ cell expression only after whole heart and basal irradiation while the apical region remained unaffected. Irradiation of the base alone led to an increase in CD45^+^ cells which was greater than that seen after whole heart or apical irradiation. This along with the fact that basal irradiation led to development of significantly greater number of lesions in the basal coronary arteries and the myocardium, indicate that the combination of age, HFD and exposure of the base of the heart to radiation, are the critical components resulting in accelerated CVD.

Platelet/endothelial cell adhesion molecule-1 (PECAM-1) or CD31, is an endothelial cell surface differentiation antigen, expressed on the surface of human granulocytes, monocytes and platelets [46]. PECAM-1 performs a key role in maintaining the integrity of the endothelial barrier, leukocyte trafficking and mechano-transduction. It has been demonstrated that although PECAM-1 deficient mice do not show any baseline vascular abnormalities, they respond poorly to inflammatory challenges owing to lack of PECAM-1 at endothelial cell-cell junctions [47, 48]. In studies designed to understand PECAM-1 expression on lung EC after radiation exposure and its impact on platelet or leukocyte-EC interactions, it has been observed that PECAM-1 has an impact on sustained functional RT-induced prothrombotic properties of EC owing to enhanced platelet-EC interactions [49]. Therefore, the RT-induced increased expression of adhesion molecules such as PECAM-1 is associated with a long-term inflammatory effect that may play a role in the development of delayed RT-induced CVD.

In our studies we observed that the increase in PECAM-1 at 8 weeks post-RT was localized to the site of irradiation. Nine weeks old mice showed increased in PECAM-1 expression in the base and apex upon whole heart RT, in the basal area upon receiving radiation at the base of the heart and in the apical region post-RT of the apex. Interestingly, older (16 weeks old) mice showed a trend towards increased PECAM-1 expression in both the base and apex regions after whole or partial cardiac irradiation and showed an increase in the baseline PECAM-1 expression as compared to the 9 week old ApoE ^-/-^ mice.

In our studies with ApoE ^-/-^ mice on HFD, the increase in PECAM-1/CD31 expression is associated with: (1) significant increases in number of lesions post-RT compared to unirradiated controls; (2) greater number of atherosclerotic lesions as compared to the other studies. These results reflect that the increased expression of adhesion molecule PECAM-1/CD31 is in fact associated with prolonged inflammation and thus accelerated development of atherosclerotic lesions.

In our studies the 16 weeks old mice on HFD exposed to cardiac RT showed no significant differences in cardiac functional parameters such as ejection fraction, stroke volume, cardiac output and fractional shortening in the mice receiving no, whole or partial cardiac irradiation assessed by ultrasound at 4 and 8 weeks post RT. However, this is not surprising since ten to eleven weeks old ApoE^-/-^ mice on a normal diet subjected to coronary occlusion with induction of myocardial infarction have been shown to have normal cardiac output, fractional shortening, and other cardiac functional parameters comparable to control age-matched C57Bl/6 mice on a normal diet subjected to coronary occlusion with induction of myocardial infarction [50, 51].

Interestingly, we observed that all mice on HFD showed the presence of multifocal perivascular mononuclear infiltrates (minimal to moderate) in the base and apex whether or not they received cardiac RT (data not shown). The presence of these infiltrates characterizes an inflammatory response and impaired vascular function. Perivascular immune cell infiltration has been shown to have a pro-atherosclerotic role associated with early stages of atherosclerotic plaque development [52]. However, whether or not these infiltrates observed during our studies indicate a process of initiation and development of atherosclerotic lesion formation remains a matter of further study.

In summary our studies demonstrate that local radiation to the heart in ApoE^-/-^ mice results in increased formation of atherosclerotic plaques and the heart base is more sensitive to these effects as compared to other areas. These findings highlight the need to avoid heart irradiation during RT as it may result in increased cardiovascular morbidity and mortality.

## Acknowledgements

Dr. Suchit Patel for assisting in establishing cardiac irradiation protocol. Dr. Liyang Zhao for experimental/technical assistance. Molecular Cytology Core facility (Yevgeniy Romin) for help with en face aorta imaging. MSKCC Small animal imaging facility (Valerie Longo) for training with cardiac ultrasound.

## Sources of funding

This work was supported in part by the Department of Radiation Oncology at MSKCC (A H-F), NIH (RO1 CA215719-04 to A.H-F and RO1 DK114321 to EAJ) and MSKCC Cancer Center Support Grant from NIH (P30 CA008748). IJK and Y.Q. are supported by NIDDK P60DK020541 (Einstein DRTC).

## Conflict of interest statement

E. A. Jaimes: Stock Shareholder: Significant; Goldilocks Therapeutics, Inc.; A. Rimner: Research Grants: Varian Medical Systems, Boehringer Ingelheim, Pfizer, AstraZeneca, Merck Consulting: AstraZeneca, Merck, Cybrexa, MoreHealth, Research To Practice.

Other authors: None

